# An invasive plant species enhances biodiversity in overgrazed pastures but inhibits its recovery in protected areas

**DOI:** 10.1101/2020.08.16.227066

**Authors:** Gianalberto Losapio, Consuelo M. De Moraes, Rodolfo Dirzo, Lilian L. Dutoit, Thomas Tscheulin, Nikos Zouros, Mark C. Mescher

**Affiliations:** Department of Environmental Systems Science, ETH Zürich, ZH 8092, Switzerland; Department of Biology, Stanford University, Stanford 94305, USA; Palo Alto, 94301, USA; Department of Geography, University of the Aegean, Mytilene 81100, Greece

**Keywords:** biodiversity-ecosystem function relationships, competition, environmental change, disturbance, indirect facilitation, plant communities

## Abstract

Anthropogenic environmental change exposes biological communities to concurrent stressors (e.g., changes in climate and land-use, overexploitation, biotic invasions) that frequently persist over prolonged periods. Predicting and mitigating the consequences of human action on nature therefore requires understanding how exposure to multiple interacting stressors alters biological communities over relevant (e.g., multi-decadal) time periods. Here, we explore the effects of overgrazing and plant species invasion on plant community diversity and ecosystem functioning (productivity), as well as the patterns of recovery of plant communities following cessation of grazing pressure. In a Mediterranean pasture system, we utilized a “natural” experiment involving long-term exclusion of grazers (for 15-25 years in parks) and also conducted short-term grazing-exclusion and invasive species removal experiments. Our results reveal that invasion by a grazing-resistant plant (prickly burnet) has net positive effects on plant diversity under overgrazing conditions but inhibits the recovery of biodiversity once grazing ceases. Furthermore, while the diversity-productivity relationship was found to be positive in pastures, the interactive effects of overgrazing and species invasion appear to disrupt ecosystem functioning and inhibit the recovery of pasture productivity. These findings highlight the potential for prolonged exposure to anthropogenic stressors, such as overgrazing, to cause potentially-irreversible changes in biological communities that, in turn, compromise ecosystem functioning and resilience. In such cases, sustainable ecosystem management may require direct intervention to boost biodiversity resilience against centennial overgrazing.

---

Human activities are causing rapid environmental change at local and global scales (1–3), leading to dramatic increases in the frequency and intensity of ecological disturbance (4,5). As a consequence, global biodiversity is rapidly declining, with potentially profound implications for critical ecosystem functions (6,7) and ecological sustainability (8–10). A key feature of anthropogenic environmental disturbance is the coincident occurrence of multiple stressors that persist over long periods of time, for example when changes in land use are accompanied by the introduction of novel species (4) or loss of native ones (10). Furthermore, the interaction between such stressors can have serious implications for important ecosystem functions, such as productivity, as well as the ability of natural systems to tolerate or recover from disturbance events (7,11–13). To predict and mitigate the long-term effects of anthropogenic environmental change on ecological sustainability, it is therefore crucial to understand how biological communities are altered by, and recover from, simultaneous exposure to multiple perturbations (3,6).

The ability of an ecosystem to recover from disturbance is termed resilience (14). In the face of anthropogenic environmental change, ecosystem resilience can be strongly influenced by changes in biodiversity and ecosystem functioning that arise from the interaction of multiple stressors acting across different spatial and temporal scales (3, 10,15). For instance, local-scale perturbations such as fire exacerbate the negative effects of large-scale deforestation (12). However, we currently have limited understanding of the processes that determine whether, and how, biological communities recover from prolonged exposure to multiple stressors or shift to a new state. Improved understanding of these processes is necessary for managing, conserving and restoring ecosystems.

In the European Mediterranean Basin, severe overgrazing across centuries of human habitation has dramatically altered ecosystem composition, structure, richness, and productivity (16–18). While some studies suggest that moderate grazing may be beneficial for the maintenance of some plant species (19,20), overgrazing is currently pushing ecosystems towards desertification in many areas, including on the Greek island of Lesvos, where this study was conducted (16). Heavily overgrazed pasture lands in Lesvos have also experienced widespread invasion by prickly burnet (*Sarcopoterium spinosum* (L.) Spach, Rosaceae), a thorny dwarf-shrub that, because of its unpalatability and high grazing resistance, has come to dominate vast areas of rangeland throughout the eastern Mediterranean Basin (18,21) (Fig. 1a and *SI Appendix,* Figure S1). This invasion has made many pastures on Lesvos nearly unusable, causing severe problems for the local human population, which rely heavily on livestock grazing for their livelihood and food production.

**Fig. 1.**
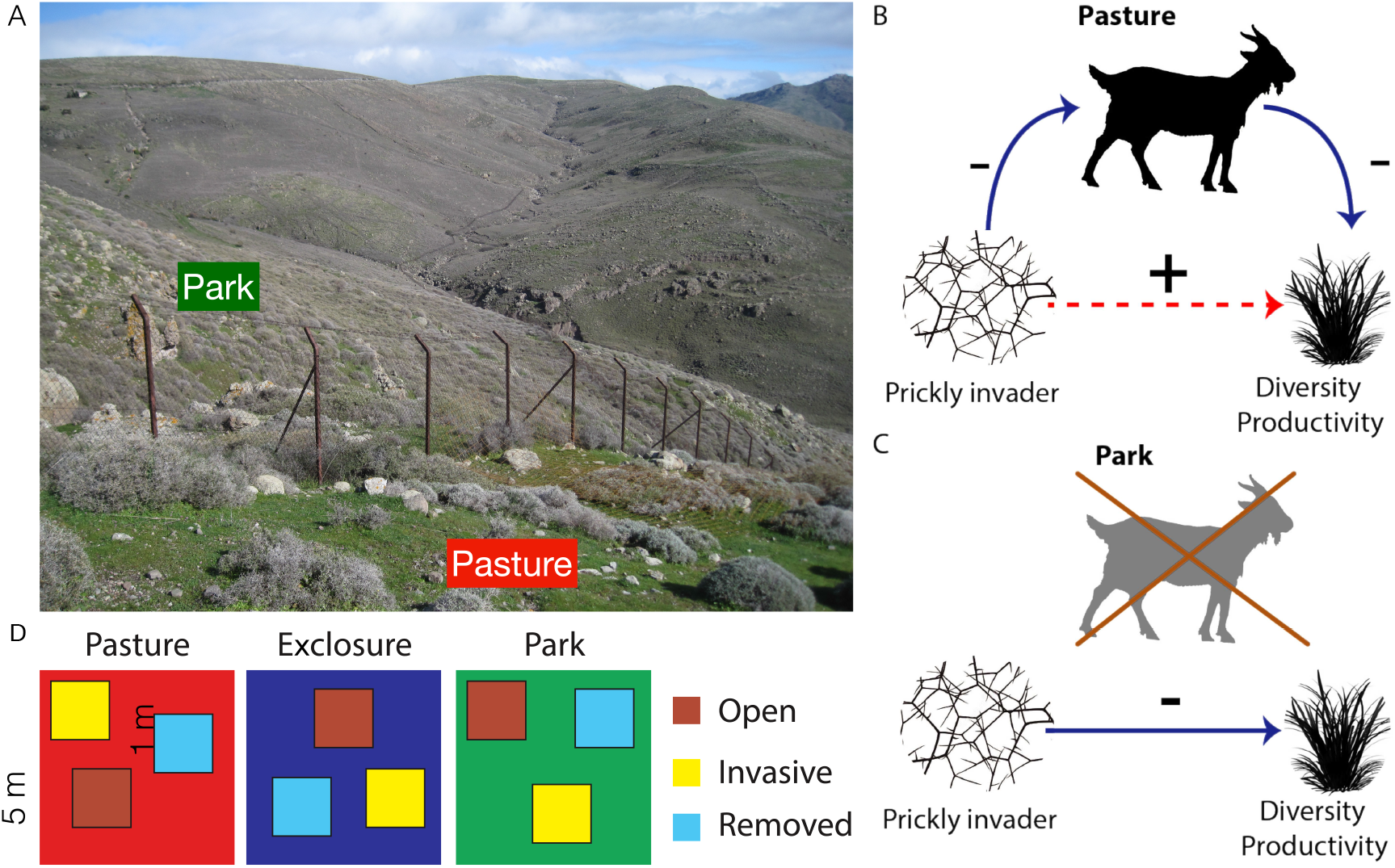
(A) Photograph of pasture and adjacent park land, separated by a fence, both invaded by prickly burnet (the greyish shrub). (B) Under overgrazing, prickly burnet protects otherwise vulnerable plants, ultimately increasing diversity. (C) Diversity-prickly burnet interactions become negative once overgrazing ceases inside exclosures and parks, limiting the resilience of biodiversity. (D) Scheme of the experimental design, comprising plots (1 m x 1 m) of species invasion treatments within blocks (5 m x 5 m) in each land management type (pasture, exclosure, park).

By reducing grazing pressure, however, prickly burnet invasion may prevent further ecosystem degradation and desertification. Indeed, there is some indication that prickly burnet can promote plant diversity by improving soil conditions and providing shelter against livestock grazing to other plants that grow beneath its thorny, protective canopy (22). Such facilitative interactions are common to pasture ecosystems, worldwide, which are frequently invaded by single, highly resistant species (23,24). However, less is known about the consequences of species invasion for ecosystem recovery once livestock grazing is halted and pastures are abandoned or set aside to become protected areas (e.g., national parks). Despite their importance, these relationships are frequently not considered in conservation programs (21).

To address whether and how species invasion mediates the impact of land-use change and overgrazing on ecosystem resilience, we studied a pasture ecosystem and adjacent protected parks on Lesvos (see Methods). Our overarching goals were to examine the long-term effects of combined overgrazing and species invasion on plant diversity and productivity and the resilience of local plant communities in the face of these stressors. Lesvos offers a unique opportunity to address these questions, since the establishment of two parks in 1994 and 2002 (UNESCO Lesvos Petrified Forest Parks in Sigri and Plaka) created a “natural” long-term exclosure experiment in which grazing was abruptly terminated on fifteen hectares of pasture lands, while intense grazing continued on adjacent lands that were otherwise similar. In addition to taking advantage of this long-term exclusion at the landscape scale, we implemented short-term exclosure and invasive-species removal experiments at the local community scale (Fig. 1, see Methods).

To assess the recovery of plant communities once overgrazing ceased, in 2018 we laid out 5 x 5 m experimental blocks for plant community surveys in: (i) grazed pastures, (ii) adjacent parks where grazing had ceased 16 or 24 years before, and (iii) fenced enclosures within the pastures. We will henceforth refer to these different areas as “land management types”. To test the consequences of species invasion and its interaction with land management, we performed a removal experiment in each block with the following treatments implemented in 1 x 1 m plots: (a) prickly burnet removed by clipping (‘removal’), (b) open areas without prickly burnet (‘open’), and (c) prickly burnet left intact (‘invasive’). Our experimental blocks were replicated twelve times at two different sites for a total of 108 1 x 1 m plots. At the end of the growing season, we measured the composition, richness, and productivity of the plant communities, which mainly comprised annual plants.

## Results and discussion

### Interactive effects of species invasion and overgrazing at landscape scale

We addressed the effects of species invasion on biodiversity resilience against overgrazing at the landscape scale. To do so, after checking for sampling completeness (*SI Appendix,* Figure S2) and standardizing by sample-based extrapolation (25,26), we quantified the effect of prickly burnet on plant diversity (*γ*-diversity) by means of a relative interaction index (27) (see Methods). When comparing plant communities in ‘invasive’ and ‘open’ areas, we found that the impact of species invasion on plant diversity differed between land management types (*SI Appendix,* Table S1). Specifically, the presence of the invader had positive effects on plant diversity in pastures but negative effects inside exclosures and parks (Fig. 2 and *SI Appendix,* Table S2). This pattern highlights the potential for one environmental stressor to revert the effects of another on ecological communities in a context-dependent way. In this case, an invasive species protects and facilitates otherwise vulnerable plants experiencing overgrazing, but also inhibits the recovery of biodiversity once such disturbance ceases.

**Fig. 2.**
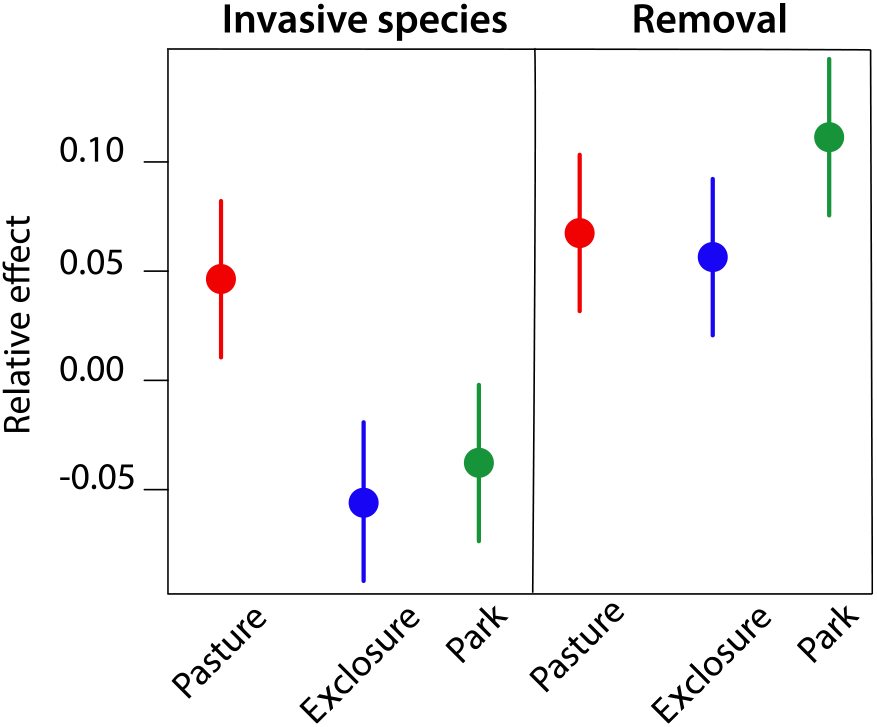
Effects of an invasive plant (prickly burnet) and its removal on plant diversity (species pool) in pasture (red), exclosure (blue) and park (green) lands. Invasive species effect is relative to open communities (without prickly burnet). Removal effect is relative to communities with prickly burnet. Dots represent estimated (marginal) means with 95% CI of the Relative Interaction Index.

Next, we explored a potential mechanism for the idiosyncratic effects of invasion on diversity described above by experimentally removing the invader. When comparing plant communities in ‘removal’ and ‘invasive’ areas, we found that removing prickly burnet had consistently positive effects on plant diversity (Fig. 2; *SI Appendix,* Table S2), indicative of a net negative (competitive) effect of prickly burnet on associated plants. Furthermore, these findings suggest a legacy effect (22,28), in which the prior presence of prickly burnet has a constructive impact on soil conditions as compared with those of open, grazed communities.

Together with our previous findings, these results suggest that species invasion enhances biodiversity maintenance in the presence of livestock but inhibits its resilience once such pressure ceases. A possible underlying mechanism is related to the shift in the balance of facilitation and competition between shrub species and associated plants (29), where net positive and negative effects prevail in pasture and in parks, respectively (Fig. 1). A similar pattern has been observed in overgrazed plant communities in Caucasus mountain ecosystems (23), suggesting that the facilitative role of unpalatable invaders for biodiversity maintenance is more relevant and widespread than previously thought.

### Persistent effects of grazing and species invasion on plant diversity and turnover

When we assessed the resilience of local-scale diversity (*α*-diversity) and community composition (*β*-diversity) against the combination of grazing and invasive species, we found that overgrazing significantly decreased biodiversity; moreover, in the absence of prickly burnet, plant communities could recover locally once grazing ceased (*SI Appendix,* Figure S3a and Table S3). In particular, local diversity increased inside exclosures (i.e., after short-term livestock exclusion from pasture lands) and increased even further in parks (i.e., after long-term livestock exclusion; *p* < 0.001).

The presence of prickly burnet also had strong effects on diversity: in pasture lands, local diversity was higher in the presence of prickly burnet and after its removal in comparison to open communities, whereas diversity was similar across these treatments inside both exclosures and parks (*SI Appendix,* Table S4). Consistent with our previous results, these findings further suggest that the positive effects of the invader on local plant diversity occur only in the presence of livestock, whereas the resilience of biodiversity may be limited by competitive effects that prevail once overgrazing disturbance ceases. Considering bottom-up and top-down drivers of plant diversity (24), our results suggest that the resilience of plant communities is regulated by both processes.

While community composition was similar in both prickly burnet and open communities across land management types, removal of the invasive species increased species turnover in parks (*SI Appendix,* Fig. S3b and Table S5). These results indicate that the presence of both livestock and invader tend to promote homogeneous plant communities, while long-term grazing exclusion combined with invader removal leads to increased community heterogeneity.

Contrary to previous claims regarding the positive effects of livestock grazing on biodiversity (16,20) but consistent with current evidence on the impact of such land-use practice (4,9,12), our experiment documented detrimental impacts of overgrazing on plant communities. Livestock grazing erodes biodiversity by favoring a few dominant species (15,23,24) and triggering invasion by unpalatable plants that ultimately limit the ability of communities to recover. Consequently, we foresee that effective management measures to improve ecosystem resilience in this and similar systems will need to include both the reduction of grazing pressure and exclusion of the invasive species.

### Grazing and species invasion jointly alter the biodiversity-ecosystem functioning (BEF) relationship

Having found positive effects of prickly burnet on plant diversity in the presence of livestock, but negative effects in its absence, we examined the combined effects of the invader and grazing pressure on ecosystem functioning. Here, we found that the resilience of productivity against the invasive species varied with land management (Fig. 3 and *SI Appendix,* Table S6). In particular, livestock grazing significantly decreased productivity, which recovered with long-term livestock exclusion in parks (*SI Appendix,* Table S7). Notably, productivity was similar between pastures and exclosures, indicating that it was independent of biomass removal (Fig. 3 and *SI Appendix,* Table S7). Furthermore, the presence of prickly burnet decreased productivity, while its removal had positive effects, particularly in parks (*SI Appendix,* Table S7). These results indicate that putting pastures under protection and removing invasive species increases productivity.

**Fig. 3.**
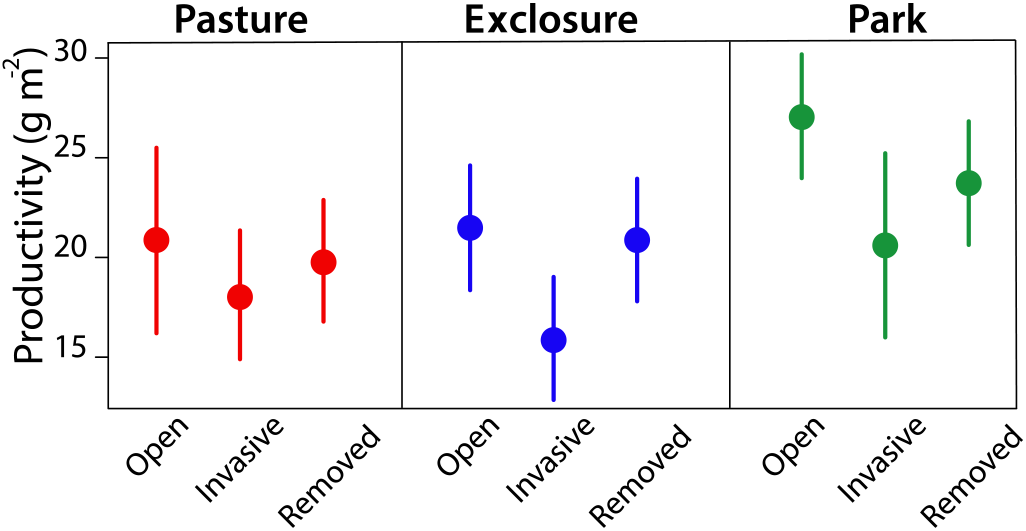
Effects of land management (i.e., pasture, exclosure and park) on productivity (g m^-2^) across treatments (i.e., open, invasive species, and its removal). Dots represent estimated (marginal) means with 95% CI.

Finally, we explored how the biodiversity-productivity relationship (3,6,11) changed under the combined impact of species invasion and land management (see Methods). Although pasture productivity generally increased with increasing plant diversity (*SI Appendix,* Table S6), the nature of this BEF relationship differed between removal treatments depending on land management (Fig. 4). Specifically, the BEF relationship was negative in the presence of prickly burnet, while it became more positive with the removal of prickly burnet. Furthermore, this trend was consistent across land management types (*SI Appendix,* Table S7 and Table S8).

**Fig. 4.**
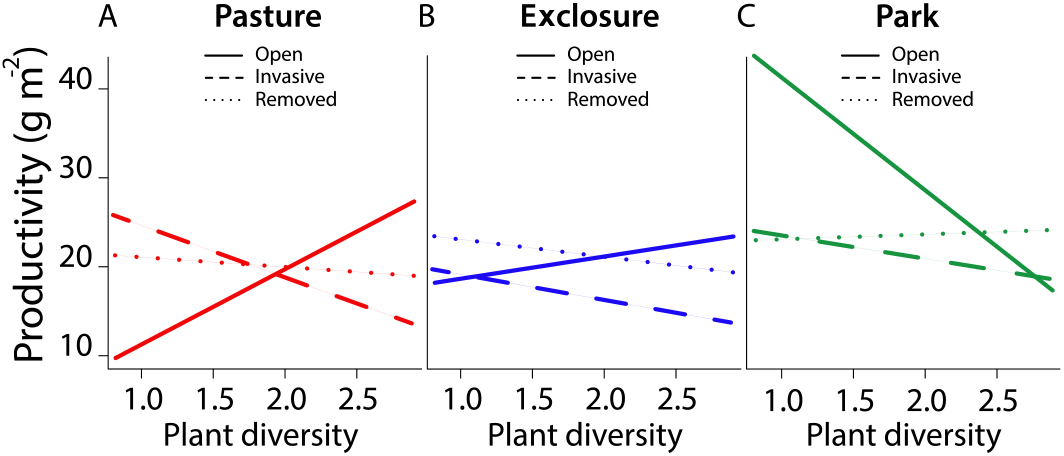
Relationships between plant diversity and productivity across land-use management and treatments. (*A*) Pasture. (*B)* Exclosure. (*C*) Park. Solid lines represent open communities; dashed lines represent invasive species communities; dotted lines represent communities where invasive species was removed.

This finding was most likely due to the previously discussed legacy effect and partial recovery, indicating that species invasion erodes the positive effects of biodiversity on ecosystem functioning. In contrast, in the absence of prickly burnet, productivity increased with plant diversity in pastures but decreased in parks. This pattern might be related to nutrient input by livestock (20), which may help maintain a positive BEF relationship. Taken together, these results indicate that changes in land management systematically alter the relationship between biodiversity and ecosystem functioning, with species invasion mediating the magnitude and sign of the diversity-productivity relationship.

### Implications for the sustainable management of Mediterranean pastures and parks

Three decades of biodiversity research suggest that more diverse ecosystems are more productive and may be more resilient to environmental perturbations through redundancy of functions carried out by different species (3,7). Furthermore, recent theoretical studies of plant networks predict that certain foundation species are critical for maintaining biodiversity as they increase community robustness (15,29). Yet, despite these advances, our understanding of the ways in which biodiversity concurrently contributes to the ability of ecosystems to recover functioning after prolonged exposure to multiple anthropogenic stressors remains far from complete. Biodiversity experiments show that greater numbers of species within communities are associated with lower invasibility (9) and higher productivity (7). Here, we show that conserving biodiversity alone may not be sufficient to ensure the long-term recovery of pasture productivity even 15-25 years after the cessation of overgrazing. This is likely due to disruption of the positive relationship between biodiversity and ecosystem functioning by species invasion.

A striking feature of our study system is the extent to which prickly burnet continues to dominate the plant community in parks, where grazing ceased decades ago. A possible explanation for this dominance is the total absence of any trees and shrubs in the landscape, which, once established, might well exclude prickly burnet from the community by limiting its recruitment. Indeed, it has been observed that prickly burnet germinates only in the absence of an overhead canopy (18,21). The observation that simply placing former pasture areas under protection is insufficient strongly suggests that proactive measures of conservation and restoration are essential (6,10).

Restoring native shrubs and trees might be an effective strategy for suppressing species invasion and boosting ecosystem resilience. Since grazing impacts ecosystem functioning and the associated canopy removal increases soil erosion and water loss (4), tree and shrub restoration may help recover multiple ecosystem functions such as productivity, soil erosion control and water retention (30). Restoring shrublands and woodlands with diverse species would also boost ecosystem resilience through the re-establishment of a positive BEF relationship. Outside parks, this measure would likely also enhance the resilience and productivity of pastures, since a drastic reduction in livestock overgrazing is needed to halt and reverse the process of desertification.

In sum, our results provide evidence that biodiversity and ecosystem functioning have limited resilience against multiple anthropogenic stressors in overgrazed landscapes. In the presence of livestock, we found that an invasive plant species helps reduce the severity of biodiversity loss at both local and landscape scales by protecting plants under its canopy. In contrast, once grazing ceases, the same invasive species inhibits biodiversity recovery, indicating potentially-irreversible biodiversity loss in the absence of targeted restoration efforts. These findings indicate that creating parks without proactive intervention is necessary but insufficient to restore desirable levels of biodiversity and ecosystem functioning due to the long-lasting detrimental effects of multiple stressors. Rather, interventions such as the restoration of native shrub and tree species may be needed to mitigate anthropogenic impact, enhance resilience and restore desirable ecosystem functioning.

## Materials and Methods

### Field experiment

The study was carried out in an overgrazed ecosystem in the western part of Lesvos Island, Greece. In this area, intense long-term grazing pressure of livestock (mainly sheep and goat) has led to domination of the plant community by a spiny invasive shrub, *Sarcopoterium spinosum* (L.), ‘prickly burnet’. *S. spinosum* is a thorny, unpalatable dwarf-shrub highly resistant to grazing (Fig. 1). It is native to the Southeast Mediterranean and Middle East (17,18,21). The establishment in some areas of UNESCO Lesvos Petrified Forest Parks in 1994 (Sigri Park) and 2002 (Plaka Park) led to the removal of overgrazing disturbance and maintenance of the ecosystem in a “natural” state. Nevertheless, prickly burnet continues to dominate the plant community inside the parks as of 2018 (Fig. 1). This system in itself constitutes a “natural” experiment allowing us to address whether and how biodiversity was able to recover from long-term overgrazing.

We therefore chose two adjacent communities differing in land-use management: pasture lands where overgrazing is still ongoing, and the above-mentioned parks where overgrazing ceased 15-25 years previously. A third land type was created with the localized exclusion of grazing in pasture lands. By excluding livestock, we could assess the direct short-term recovery of biodiversity and ecosystem functioning from overgrazing. In pastures, we randomly installed 5 x 5 m fences (1.5 m height) and selected adjacent 5 x 5 m areas. In parks, we randomly selected 5 x 5 m areas. For each land type, eight blocks were established at the Sigri Park site and four at the Plaka Park site, for a total of 36 blocks.

The field experimental manipulation also included the removal of prickly burnet. By removing prickly burnet, we could assess the effects of species invasion on the resilience of biodiversity and ecosystem functioning. The following three treatments were conducted: i) community without prickly burnet (open areas); ii) community with prickly burnet; and iii) removal of prickly burnet by clipping its canopy. We followed a fully-factorial, randomized block design by replicating and making blocks of the three treatments together (i.e., open areas, prickly burnet and prickly burnet removal) in each land management type (i.e., pasture, exclosure and park) (Fig. 1).

The treatment plots measured 1 x 1 m. In total, we installed *n* = 108 plots at the start of the vegetative season (February 2018). Then, we visually recorded the occurrence and cover of plant species within each plot (April 2018), using the nomenclature of Bazos (2005). Finally, we harvested the aboveground biomass of each entire plot at the end of the season (May 2018). Biomass was dried for 72 h at 70°C.

### Data analysis

First, we measured biodiversity at the landscape scale, i.e., gamma (*γ*) diversity. This measure of biodiversity refers to the species richness in a landscape, i.e., species pool. In order to cover the highest possible extent of biodiversity, we quantified the species pool by integrating the observed species richness with the estimated number of unseen (missing) species in each treatment. We used the bootstrap estimator 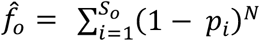 and resampled 99 times from a normal distribution with mean 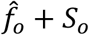 and standard deviation 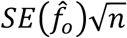 using the *specpool* function of the *vegan* R package (26,31).

We then analyzed the interactions of land management and species invasion with biodiversity. To ensure we included all plant species in the ecosystem (i.e., both observed and potentially unseen species) we used the estimated *γ*-diversity for this analysis. This also allowed us to look at biodiversity resilience at the landscape scale. We considered the following scenarios: 1) grazing and invader; 2) grazing without invader; 3) short-term grazing exclusion with invader; 4) short-term grazing exclusion without invader; 5) long-term grazing exclusion with invader; 6) long-term grazing exclusion without invader. We measured interactions between prickly burnet and *γ*-diversity using the Relative Interaction Index (RII) (27). This RII was calculated as (*γ intact – γ absent*)/(*γ intact + γ absent*) and (*γ intact – γ removed*)/(*γ intact + γ removed*) to assess the effects of prickly burnet presence and prickly burnet removal in each combination of grazing treatment, respectively. In scenarios 1 and 2 we considered pastures, in scenarios 3 and 4 we considered exclosures, in scenarios 5 and 6 we considered park lands. RII ranges from −1 to 1, with values smaller and higher than 0 indicating negative and positive interactions between plants and prickly burnet, respectively. RII was tested in response to grazing (presence/exclusion) and invader (presence/removal) with time scale (short- and long-term effect) nested within grazing exclusion using hierarchical models (32); replicate pairs were random effects. We ran comparison between specific levels using least-square mean estimation (33).

We also quantified alpha (*α*) and beta (*β*) diversity. *α*-diversity refers to the observed local diversity in each plot. It was quantified using the Shannon diversity index as 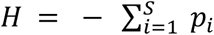 In *p_i_*, where *p* is the relative cover of plant species *i* occurring in the community with *S* species. This index accounts for both abundance and evenness of plant species. *β*-diversity refers to the dissimilarity (heterogeneity) in species composition among communities, and indicates how many different and unique species there are between each pair of plots. This was quantified using the Sørensen similarity index (26,33) as *β*-diversity = *b* + /(*2a+b+c*), where *a* is the number of shared species between two plots, *b* and *c* are the numbers of species unique to each plot.

We used hierarchical models (32,34) to test for differences among species invasion treatments (open, prickly burnet, and removal) across land management types (pasture, exclosure, and park), and their statistical interaction for indices of *α*- and *β*-diversity. Site (parks) and block nested within site were considered as random effects. For each fitted model, the significance of predictors was tested using type II Wald chi-square test in terms of explained variance (33).

We then analyzed: (i) the response of aboveground biomass productivity (g m^-2^), a classic measure of ecosystem functioning (7), and (ii) the relationship between biodiversity and productivity (BEF) over land management and species invasion. We used a hierarchical model (32) to test for differences in productivity between land management (pasture, exclosure, and park), species invasion (open, prickly burnet, and removal), and their statistical interaction as a function of *α*-diversity. Site (parks) and block nested within site were considered as random effects. We first reported the effects of predictors considering the fitting of coefficient estimates. We then estimated the diversity-productivity relationship (BEF) across treatments and land management using marginal means of linear trends (35). Data analysis was performed in R ver. 3.5.0 (31).

## Data Availability

The data collected for this study have been deposited on the ETH Research Collection server at https://www.research-collection.ethz.ch/handle/20.500.11850/309353 (doi: 10.3929/ethz-b-000309353) and https://www.research-collection.ethz.ch/handle/20.500.11850/311948 (doi: 10.3929/ethz-b-000311948). The R code to reproduce the analyses and figures will be deposited upon acceptance of the manuscript.

## Acknowledgments

GL acknowledges support from the Swiss National Science Foundation under the Scientific Exchange program (Grant n. IZSEZ0_180195) and the Early Postdoc.mobility fellowship (Grant n. P2ZHP3_187938). We thank Volker Nickels, George Karlis and Inna Nenova for their help during fieldwork.

## Author contributions

G.L., T.T., and M.C.M. conceived the study; G.L., C.M.D.M., T.T., and M.C.M designed the research; G.L. performed research, analyzed data and wrote the paper with help from L.L.D, and M.C.M. All authors contributed to the interpretation of results and provided input on the manuscript.

The authors declare no conflict of interest.

## Supplementary Information

is available in the online version of the paper.

